# Systematic genome-scale identification of host factors for SARS-CoV-2 infection across models yields a core single gene dependency; *ACE2*

**DOI:** 10.1101/2021.06.28.450244

**Authors:** Katherine Chan, Adrian Granda Farias, Hunsang Lee, Furkan Guvenc, Patricia Mero, Kamaldeep Aulakh, Kevin R. Brown, Shahan Haider, Edyta Marcon, Ulrich Braunschweig, Amy Hin Yan Tong, Shuye Pu, Andrea Habsid, Natasha Christie-Holmes, Patrick Budylowski, Audrey Astori, Ayoob Ghalami, Samira Mubareka, Arinjay Banerjee, Karen L. Mossman, Jack Greenblatt, Scott D. Gray-Owen, Brian Raught, Benjamin J. Blencowe, Mikko Taipale, Jason Moffat

## Abstract

SARS-CoV-2, depends on host cell components for replication, therefore the identification of virus-host dependencies offers an effective way to elucidate mechanisms involved in viral infection. Such host factors may be necessary for infection and replication of SARS-CoV-2 and, if druggable, presents an attractive strategy for anti-viral therapy. We performed genome wide CRISPR knockout screens in Vero E6 cells and 4 human cell lines including Calu-3, Caco-2, Hek293 and Huh7 to identify genetic regulators of SARS-CoV-2 infection. Our findings identified only *ACE2*, the cognate SARS-CoV-2 entry receptor, as a common host dependency factor across all cell lines, while all other host genes identified were cell line specific including known factors *TMPRSS2* and *CTSL*. Several of the discovered host-dependency factors converged on pathways involved in cell signalling, lipid metabolism, immune pathways and chromatin modulation. Notably, chromatin modulator genes *KMT2C* and *KDM6A* in Calu-3 cells had the strongest impact in preventing SARS-CoV-2 infection when perturbed. Overall, the network of host factors that have been identified will be broadly applicable to understanding the impact of SARS-CoV-2 on human cells and facilitate the development of host-directed therapies.

**IN BRIEF:** SARS-CoV-2, depends on host cell components for infection and replication. Genome-wide CRISPR screens were performed in multiple human cell lines to elucidate common host dependencies required for SARS-CoV-2 infection. Only ACE2, the cognate SARS-CoV-2 entry receptor, was common amongst cell lines, while all other host genes identified were cell line specific, several of which converged on pathways involved in cell signalling, lipid metabolism, immune pathways, and chromatin modulation. Overall, a network of host factors was identified that will be broadly applicable to understanding the impact of SARS-CoV-2 on human cells and facilitate productive targeting of host genes and pathways.

**HIGHLIGHTS:**

- Genome-wide CRISPR screens for SARS-CoV-2 in multiple human cell lines
- Identification of wide-ranging cell-type dependent genetic dependencies for SARS-CoV-2 infection
- ACE2 is the only common host factor identified across different cell types

## INTRODUCTION

Since its identification in late 2019, the highly infectious severe acute respiratory syndrome coronavirus (SARS-CoV-2) has rapidly spread across the world, resulting in over 160 million infections and greater than 3.7 million deaths as of June 9, 2021 (https://www.worldometers.info/coronavirus/). SARS-CoV-2 is closely related to SARS-CoV-1, which caused the 2002-2004 SARS epidemic, as well as MERS-CoV, the virus responsible for respiratory disease outbreaks in 27 countries since 2012 (da Costa et al., 2020). Infections can range from asymptomatic or mild cases to severe respiratory disease, that can lead to acute respiratory distress syndrome, septic shock and multi-organ failure (Vos et al., 2021). SARS-CoV-2, after SARS-CoV and MERS-CoV, is the third zoonotic coronavirus in the last two decades to rise to epidemic, and now, pandemic proportions. Each of these viral outbreaks can be traced to animal hosts, with zoonosis accounting for human CoV infections. Increasingly close contact with diverse animal species will continue to drive zoonotic transmission of potentially deadly coronaviruses, which underscores the importance of clearly delineating the molecular mechanisms of human CoV viral infection (Banerjee et al., 2021a; Banerjee et al., 2021c).

Like all viruses, SARS-CoV-2 is an obligate intracellular pathogen that depends on host cell components for replication. The viral envelope is studded with several proteins, including the Spike (S) glycoprotein that mediates direct virus-host cell interactions (Hartenian et al., 2020; V’Kovski et al., 2021). The SARS-CoV-2 S-protein is a heavily glycosylated homotrimer composed of two subunits, S1 (receptor binding domain - RBD) and S2 (helical domain that mediates membrane fusion), which act in a coordinate manner to attach to the host cell surface and initiate viral entry (Letko et al., 2020). S-protein RBD binds to the host receptor angiotensin-converting enzyme 2 (ACE2), which also serves as the entry receptor for SARS-CoV (Hoffmann et al., 2020). High variability in the RBD was the major determinant of cross species transmission and evolution of SARS-CoV-2 (Starr et al., 2020). Furthermore, mutations in RBD have appeared among pandemic variants, increasing virus transmissibility and potentially compromising efficacy of treatments and vaccines in development (Linsky et al., 2020). ACE2 is highly expressed in the lung, heart, arteries, kidney and intestines, an expression pattern that may in part explain the constellation of symptoms associated with SARS infection (Crackower et al., 2002; Danilczyk et al., 2006; Hashimoto et al., 2012; Imai et al., 2005; Zhang et al., 2020). In addition to ACE2, transmembrane protease serine 2 (TMPRSS2) has been identified as a cofactor that aids viral entry by priming S-protein through cleavage of the S1/S2 and S2’ cleavage sites (Zhu et al., 2021). Similarly, cathepsin L (CTSL) has also been shown to mediate S priming and facilitate intracellular viral release (Zhao et al., 2021). Recent studies show that SARS-CoV-2 has an insertion of a polybasic (RXYR) furin cleavage site at the S1/S2 boundary that is absent from SARS-CoV and other 2b betaCoVs. This could be an additional factor contributing to the increased virulence of SARS-CoV-2, as furin proteases are ubiquitously expressed in humans, providing expanded tissue tropism and pathogenesis (Wang et al., 2020).

The SARS-CoV-2 life cycle appears to require a mixed array of host factors. A deeper understanding of the host-pathogen interactions of SARS-CoV-2 across a range of permissive models will help shed light on viral pathogenesis and host defense mechanisms. In fact, therapeutic strategies targeting host factors are still important to consider as virus-targeted therapies will likely drive viral escape mechanisms and the emergence of novel variants that continue to infect at-risk populations.

Functional genetic screens using genome-wide pooled CRISPR-Cas9 libraries are designed to link genotype-to-phenotype relationships and are ideally suited to understand mechanisms of disease. Over the past five years, CRISPR-based genetic screens have improved our understanding of viral dependencies providing a powerful means for investigating viral life cycles (McDougall et al., 2018). A number of recent studies have employed loss-of-function CRISPR-based survival screening approaches to identify host factors that regulate SARS-CoV-2 viral entry (Baggen et al., 2021; Bailey and Diamond, 2021; Baumann, 2021; Daniloski et al., 2021; Heaton et al., 2020; Hoffmann et al., 2021; Schneider et al., 2021; Wang et al., 2021; Wei et al., 2021). Collectively, these studies have provided valuable insights into the host factors involved in SARS-CoV-2 infection. However, these studies employed host gene knockout either in non-epithelial cells lines or cell lines that do not endogenously express ACE2 and TMPRSS2 (Baggen *et al*., 2021; Daniloski *et al*., 2021; Schneider *et al*., 2021; Wang *et al*., 2021; Wei *et al*., 2021; Zhu *et al*., 2021). Nonetheless, each of the screens has focused on a single cell type and present predominantly non-overlapping sets of host factors, raising questions about the nature of the discrepancies (Bailey and Diamond, 2021).

We set out to systematically define host factors that regulate SARS-CoV-2 infection in multiple permissive models including Calu-3 cells, which are human lung cells endogenously expressing both ACE2 and TMPRSS2. We first performed tropism studies in multiple model cell lines and then conducted genome-wide loss-of-function CRISPR screens to generate a systematic map of host dependency factors. Pathway and comparative analysis with previous studies revealed diverse cellular pathways involved in modulating signalling processes to evade proteotoxic stress, lipid metabolism, immune pathways and epigenetic chromatin modulation.

Further to our screening observations, global transcriptional analysis of infected cell models and additional studies revealed unappreciated host dependencies and protease re-wiring outcomes in different models. Our results recapitulate that *ACE2* is the most prominent pro-viral host factor that is required for SARS-CoV-2 infection, while other genes modulate cell survival upon infection in a cell and context-dependent manner, likely due to inherent redundancy in cellular factors that the virus can utilize to complete its life cycle.

## RESULTS

### Tropism of SARS-CoV-2 in cell culture models

The SARS lineage (SARS-CoV-1 and SARS-CoV-2) of coronaviruses are typically isolated and grown on African green monkey *(Chlorocebus sp*.)-derived kidney epithelial cells (Vero E6) enriched for expression of ACE2. To date, human cell lines reported to support SARS-CoV-2 replication include Caco-2 colorectal adenocarcinoma cells, Calu-3 lung adenocarcinoma cells, as well as Huh7 and Huh7.5 hepatocellular carcinoma cells (Chu et al., 2020; Harcourt et al., 2020a; b). Most of the cell lines used for *in vitro* studies to date have required exogenous expression of *ACE2*, with or without exogenous expression of *TMPRSS2*. For example, the A549 lung adenocarcinoma and human embryonic kidney HEK293 cell lines are often used in conjunction with overexpression of *ACE2* for studying SARS-CoV-2 cellular infection (Gordon et al., 2020). Before starting genome-wide CRISPR screens for host-cell entry factors, we assessed the susceptibility of human cell lines originating from various tissues focusing on the cell types shown to be susceptible to SARS-CoV-1. The cell lines that were tested included Caco-2, Calu-3, Huh7, and HEK293 cells transduced with *ACE2* and *TMPRSS2* (HEK293^+A+T^)(Abe et al., 2020), as well as Vero E6. Each cell line was inoculated with SARS-CoV-2 at a multiplicity of infection (MOI) of 1 or 10 (Banerjee et al., 2020). Cells were monitored for cytopathic effects (CPE) and the degree of infection was determined by quantifying SARS-CoV-2 nucleocapsid (N) protein by immunofluorescence (IF) over a period of 24 to 120 hours post infection (hpi)(Figure 1a).

**Figure 1.**
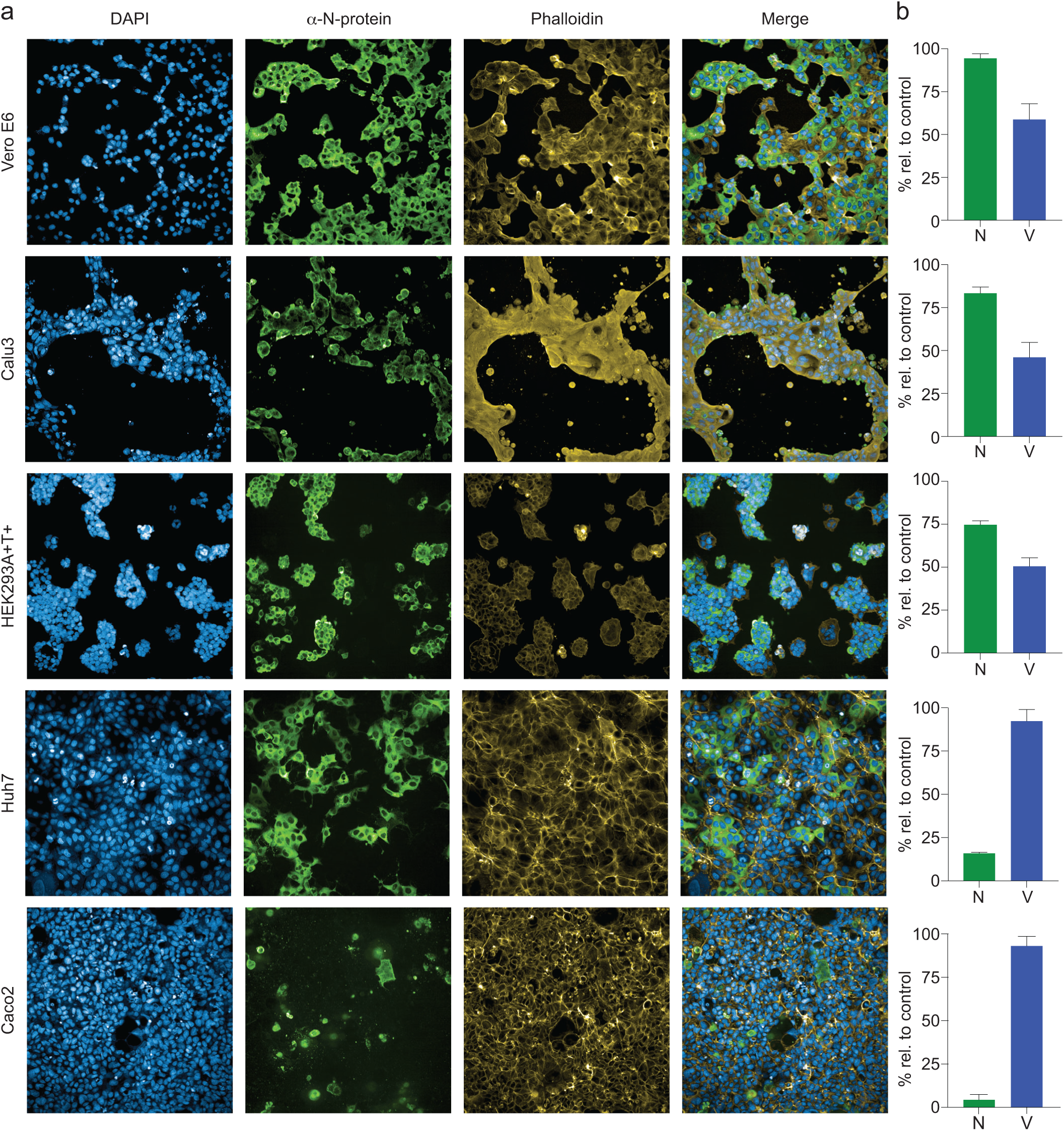
SARS-CoV-2 infection and induced cytopathic effects in human cell types SARS-CoV-2 infectivity and induced cytopathic effects were determined in Vero E6, Calu-3, HEK293^+A+T^, Huh7 and Caco-2 cells. Cells were infected with SARS-CoV-2 at MOI of 1 or 10 (Caco-2 cells). At 48 hpi, cells were fixed in 10% NBF and immunostained with anti-SARS N-protein (green), DAPI (blue) and phalloidin (yellow). A) Images were acquired with Phenix Opera system. B) The % cells positive for N-protein staining (green bars, N) was determined by quantifying number of greens cells relative to DAPI stained cells in the same well, while % cell viability was quantified by counting the number of nuclei in the infected cells relative to mock uninfected wells (blue bars, V).

Typical cytopathic effects include cell rounding, detachment, degeneration and syncytium formation, and these features were observed in Vero E6, Calu-3, and HEK293^+A+T^ cell lines coinciding with a robust infection as indicated by the high percentage of N-protein positive cells (Figure 1a,b). Huh7 and Caco-2 showed approximately 16% and 5% infection after 72 hours and 120 hours respectively, with Caco-2 cells requiring an MOI>10 to achieve any detectable infection. No significant spread to other cells occurred over the course of infection in Huh7 and Caco-2 cells, and as a result minimal CPE was observed. Representative time course of infection is shown for Calu-3 (CPE) and Huh7 (non-CPE) cells in Supplemental Figure 1. The presence of viral proteins in infected cells was measured over time by mass spectrometry and N-protein could be detected as early as 12 hours and increased over time (Supplemental Figure 2). N-protein levels were highest in Vero E6 and Calu-3 cells, with quantities increasing over time, confirming replication and production of SARS-CoV-2 in these cells. Huh7 and Caco-2 cells produced lower levels of N-protein, with Caco-2 being the lowest (Supplemental Figure 2). Spike (S) and the membrane glycoprotein (M) were also measured by mass spectrometry and the levels of both proteins increased over time in Vero E6, Calu-3 and Huh7 cells, in which infection was seen to spread in the cells (Supplemental Figure 2). However, in Caco-2 cells, S and M levels remained low and undetectable, confirming the lower transmission and replication capacity in this cell type.

An analysis of RNA expression data for each cell line suggests that Calu-3 cells have higher levels of *ACE2* expression compared to Huh7 and Caco-2 cells, suggesting that the level of *ACE2* is a key determinant of the level of infection and CPE we observed, as no CPE was detected in Huh7 and Caco-2 cells (Supplementary Table 1). In addition to transcript levels, total levels of known host entry factors ACE2, CTSL and TMPRSS2 were evaluated by western blot and flow cytometry. ACE2 was detected in all cell lines tested, with Caco-2 having comparatively much lower expression than the other cell lines (Figures 2a,b), which likely explains the infection dynamics observed in this cell line. TMPRSS2 was only detected in Calu-3 and in TMPRSS2 transduced HEK293^+A+T^ cells. In contrast, CTSL was highly expressed in Vero and Huh7 cells, but only modestly expressed in Calu-3 and Caco-2 cells (Figure 2a). Notably, protein lysates treated with the de-glycosylase PNGase showed marked shifts in ACE2 migration by Western blot analyses, indicating that ACE2 is glycosylated in these cell lines (Figure 2c).

**Figure 2.**
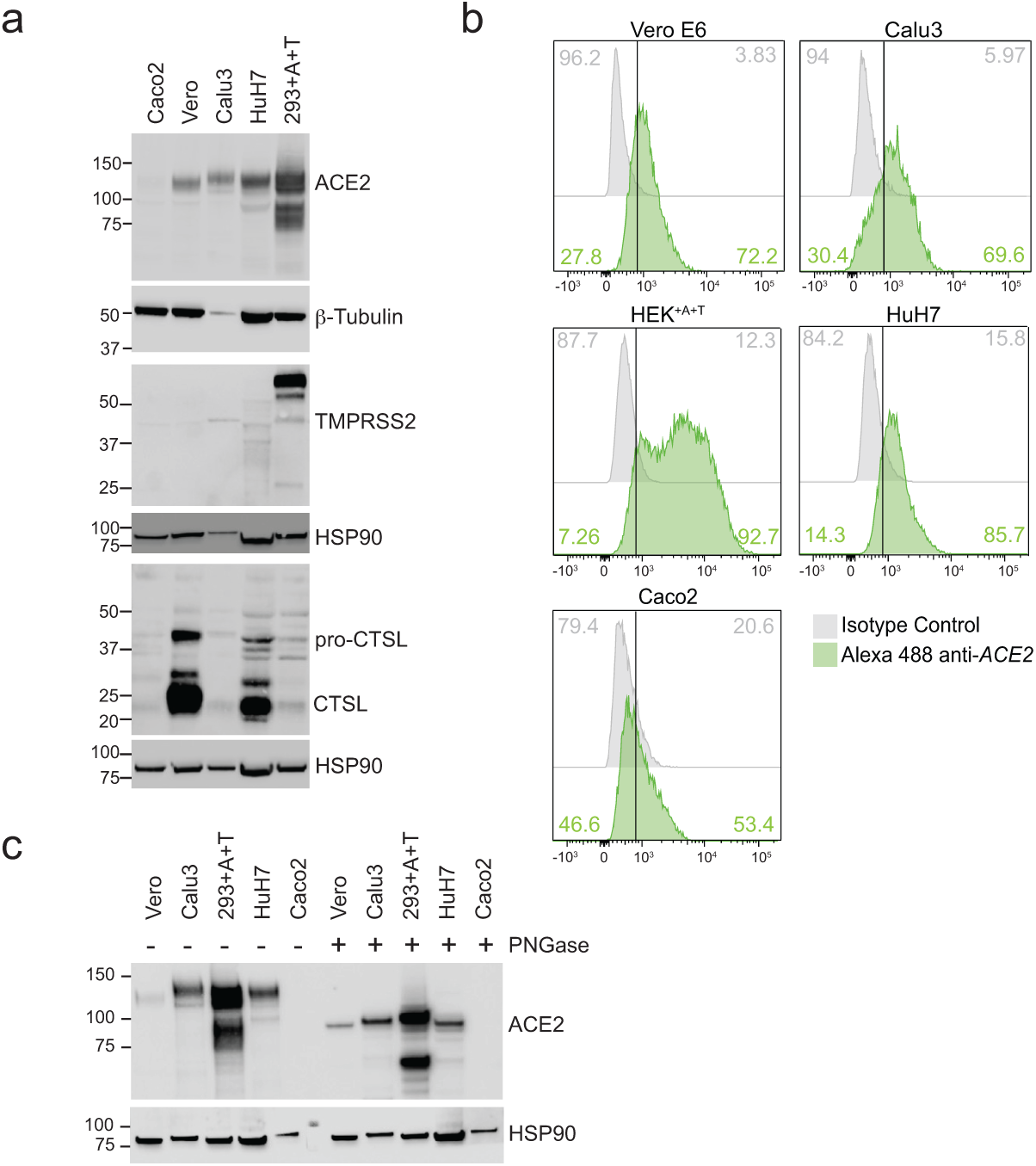
Expression levels of SARS-CoV-2 entry factors across human cell types. A) Western blot for ACE2, CTSL and TMPRSS2 expression B) Flow cytometry for ACE2 surface expression, ACE2 stained cells (green), isotype control (grey) C) Western blot for total ACE2 +/- PNGase F treatment

### Genome-wide CRISPR Screens to identify host genes required for SARS-CoV-2 cytotoxicity

To identify host genes required for SARS-CoV-2 infection we performed genome-wide CRISPR screens in Vero E6 cells and three human cell lines with different susceptibilities to SARS-CoV-2 infection including Caco-2, Huh7, and Calu-3 cells. In addition, we screened the human embryonic kidney derived HEK293^+A+T^ cell line overexpressing *ACE2* and *TMPRSS2*. As a result, three different cell types from different tissues affected by SARS-CoV-2 infections (lung, liver, and colon) were screened with the aim of identifying core genetic host factors, as well as context specific genetic factors required for SARS-CoV-2 infection and replication.

Screens were performed using the sequence optimized human all-in-one Toronto KnockOut v3 CRISPR-Cas9 library (Hart et al., 2017). The TKOv3 library contains 71,091 gRNAS (4 gRNAs per gene) targeting 18,053 protein-coding genes in the human genome (Hart *et al*., 2017). The same library was used to screen African green monkey Vero E6 cells; guides were mapped to the African green monkey reference genome to identify guides with perfect homology to their targets resulting in 56% of the TKOv3 library mapping with perfect homology and targeting 15,935 loci in the Vero E6 genome. The TKOv3 guides with no targets in the African green monkey genome also had no significant off-target matches, indicating that the non-targeting fraction of TKOv3 would not increase detection of off-target (non-specific) hits (Supplemental Table 2). Following transduction of host cells with TKOv3 lentivirus at a low multiplicity of infection (∼ MOI 0.3), cells were challenged with SARS-CoV-2 and screens were then carried out based on the presence or absence of CPE in response to infection (Figure 3a). For cells that exhibited CPE, positive selection screens based on cell-survival were performed to select for host genes that, when knocked out, confer resistance to SARS-CoV-2 infection, thus identifying pro-viral genes. Infection of Calu-3, Vero E6 and HEK293^+A+T^ pooled genome-wide knockout populations with SARS-CoV-2 resulted in >80% virus-induced cell death. The surviving cells were harvested and expanded for a second round of infection with SARS-CoV-2, after which no additional cell death was observed. Caco-2 and Huh7 cells, in which CPE was not observed, were passaged in the presence of SARS-CoV-2 for up to 15 cell population doublings. This selection condition offers the possibility of identifying gene knockouts that sensitize cells to SARS-CoV-2 infection (i.e., potential host antiviral genes). Huh7 and Caco-2 cells were infected with SARS-CoV-2 at an MOI of 1 and 10, respectively, and passaged every 3-4 days with reinfection with SARS-CoV-2 for two (Huh7) and three rounds (Caco-2). No CPE was detected at any time over the course of these screens. For all screens, genomic DNA was harvested at the final time point and guide abundance was quantified by next-generation sequencing.

**Figure 3.**
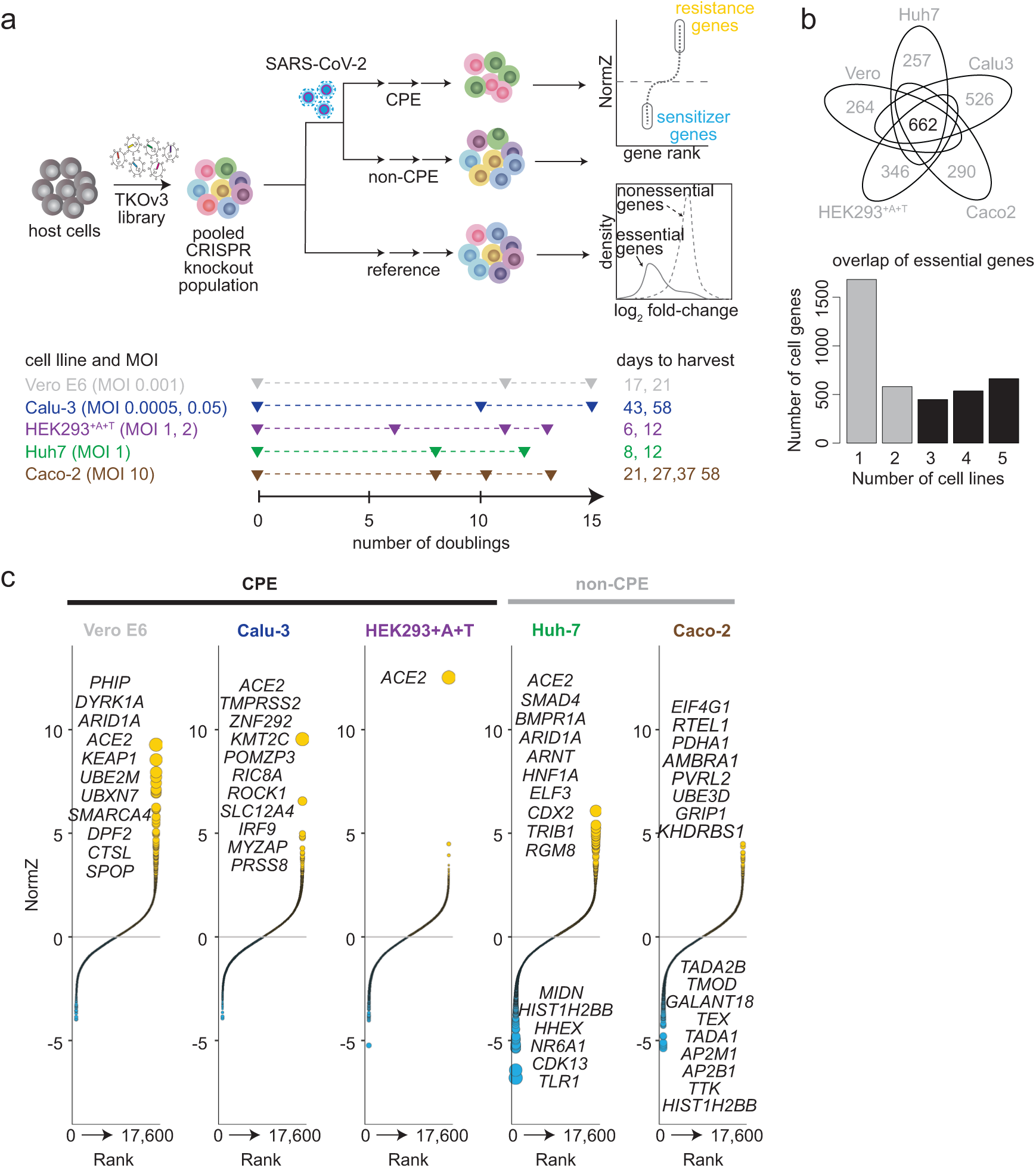
Genome-wide CRISPR screens identify host factors required for SARS-CoV-2 infection. A) Schematic of pooled genome-wide CRISPR screens and experimental timelines. Host cells were transduced with TKOv3 lentivirus. Following puromycin selection of the transduced population, cells were infected with SARS-CoV-2 or mock treated. Surviving cells were isolated and sgRNA abundance was determined next generation sequencing to identify host factors. B) Daisy model of gene essentiality across different cell lines and distribution of hits. Petals represent essential genes unique to each cell line. The bar graph represents distribution of hits at 5% FDR across the cell lines. Black font/bars represent core essentials; grey font/bars represent context essentials. C) Gene-level normZ scores showing the top genes conferring resistance (yellow) and sensitivity (blue) to SARS-CoV-2. Hits at FDR<10% are highlighted, and the dot size is inversely scaled by FDR.

The overall performance of all the screens was robust as evaluated by changes in cell proliferation or fitness when targeting gold standard reference sets of essential and non-essential genes (Hart *et al*., 2017). Precision-recall (PR) curves show that at 5% FDR, >95% of essential genes were identified, confirming efficient Cas9 editing and disruption of genes in our screens. Additionally, the fold-change distribution of gRNAs targeting essential genes was significantly shifted relative to those targeting nonessential genes, further confirming robust Cas9 editing in these populations (Supplemental Figure 3a). Pairwise correlation analyses of time point data also demonstrated the T0 time point of each genetic screen clustered together with high correlation coefficients, as did technical replicates, confirming that TKOv3 was equally represented across all screens at the onset of TKOv3 infection (Supplemental Figure 3b).

Using the Daisy model of gene essentiality (Hart et al., 2014; Hart et al., 2015; Hart and Moffat, 2016; Hart *et al*., 2017), we evaluated gene essentiality across the cell lines screened to identify core essential genes and context-specific genetic vulnerabilities. This analysis revealed 662 core essential genes that were detected in all five cell lines (BF>5, FDR <0.05) and subsets of context-dependent genes that were specific to each cell line (Figure 3b, Supplemental Table 3). The context-essential genes in each cell line were subjected to gene set enrichment analysis (GSEA), to distinguish unique signatures of essential biological processes and associated components for each cell line. Only Vero E6 hits showed unique enrichment for DNA damage response pathways (adj pval <0.05), which is consistent with its *TP53* wildtype status compared to the other cell lines which were *TP53* mutants (Brown et al., 2019) (Supplemental Figure 4a). Calu-3, Caco-2, HEK293^+A+T^ and Huh7 context-essential gene sets were consistent with genes in the amplified regions of their genome, as determined by chromosome position analysis of screening data (Supplemental Figure 4b). Biological process annotation of genes in the amplified gene regions were similar across the cell lines and consisted of vital cell processes such as RNA splicing, translation, cell cycle, and DNA replication. None of the identified gene hits discussed below were core or context essential fitness genes.

Genes within the pool when perturbed that led to enrichment or depletion as a result of SARS-CoV-2 were scored using drugZ algorithm (Colic et al., 2019)(Figure 3c, Supplemental Table 4). As expected, *ACE2* was identified as a top resistance hit in the SARS-CoV-2 screens that exhibited good infection including Vero E6, Calu-3 and HEK293^+A+T^ cells, validating the capability of our screens to select for host factors required for SARS-CoV-2 cytotoxicity. *ACE2* was also positively selected in Huh7 cells (FDR<0.2), where CPE was not observed during the screen. *ACE2* was not captured in Caco-2 cells, which we attribute to lower levels of infection achieved in these cells, indicating that the selection pressure was not robust enough to enrich for these guides. In terms of co-factors that are known to facilitate SARS-CoV-2 infection, several notable observations were made. *CTSL*, which encodes the Cathepsin L protease, was positively selected only in Vero E6 cells while the serine protease *TMPRSS2* was only positively selected in Calu-3 cells (Figures 3c). These results point out that the protease used for priming S-protein, and mode of entry, are likely cell-type dependent. Moreover, the top and only significant hit in our HEK293^+A+T^ screen was *ACE2*, suggesting that overexpression of *ACE2* is sufficient for SARS-CoV-2 infection in this cell line, and *TMPRSS2* over-expression may be important in this situation for improving infection of viral particles pseudotyped with Spike protein, but not so much for infection by authentic SARS-CoV-2 viruses. Indeed, this is consistent with a previous report, in which inhibition of cathepsin L (CTSL) and cathepsin B (CTSB) in 293T+ACE2 cells reduced entry of pseudotyped-Spike viruses, whereas inhibiting TMPRSS2 did not, indicating that SARS-CoV-2 entry into 293T-ACE2 cells depends on CTSB or CTSL. Therefore, it appears that exogenous TMPRSS2 may be redundant due to the underlying effect of endogenous CTSB/L expression, which was not identified in our HEK293^+A+T^ screen (Crawford et al., 2020; Hu et al., 2020; Ikegame et al., 2021).

### Enriched pro-viral genes reveal common mechanisms for viral hijacking of host cellular processes

Each cell line screened in the presence of SARS-CoV-2 revealed different gene hits. Closer examination of the top 10 significant hits (Figure 3c, FDR <0.1) suggest the most enriched pro-viral genes identified various scenarios for which SARS-CoV-2 interfere with signalling networks to evade host cytoprotective processes and facilitate viral pathogenesis, that have also been observed with other virial pathogens (Figure 4). The top Vero hits were genes that encode components of the Cullin3-based Cullin Ring E3 ligase family (CRL3) complex (*KEAP1*, *UBE2M*, *SPOP*, *UBXN7*) and Cullin4-based CRL complex (*PHIP*). CRLs are ubiquitination complexes that regulate viral cell processes. Notably, CUL3 complexes recruit substrates for ubiquitination through BTB-domain receptor substrate binding motifs (Petroski and Deshaies, 2005), of which are also found on several virus proteins (Mahon et al., 2014). KEAP1, a CUL3 substrate receptor, negatively regulates transcription factor NRF2, which activates cytoprotective response to environmental stress allowing cell survival under stress conditions. Viruses, like MARVs, EBOVs, HBV, HCV and CMV have been shown to disrupt the KEAP1-NRF2 interaction, activating NRF2 to induce cytoprotective responses during infection (Edwards et al., 2014). Genes encoding chromatin remodeling factors (*ARID1A*, *SMARCA4*, *DPF2*, *DYRKIA*) were also top hits. Although RNA viruses such as SARS cannot hack the host genetic sequence, recent research suggests RNA viruses can utilize aspects of the epigenetic machinery to enable infection, spread and transmission by antagonizing the immune system through epigenetic mechanisms (Atlante et al., 2020). For example, DPF2, a negative regulator of non-canonical NF-kß pathway, is used by influenza to evade the host immune system by suppressing IFN-ß expression and thus its production (Shin et al., 2017).

**Figure 4.**
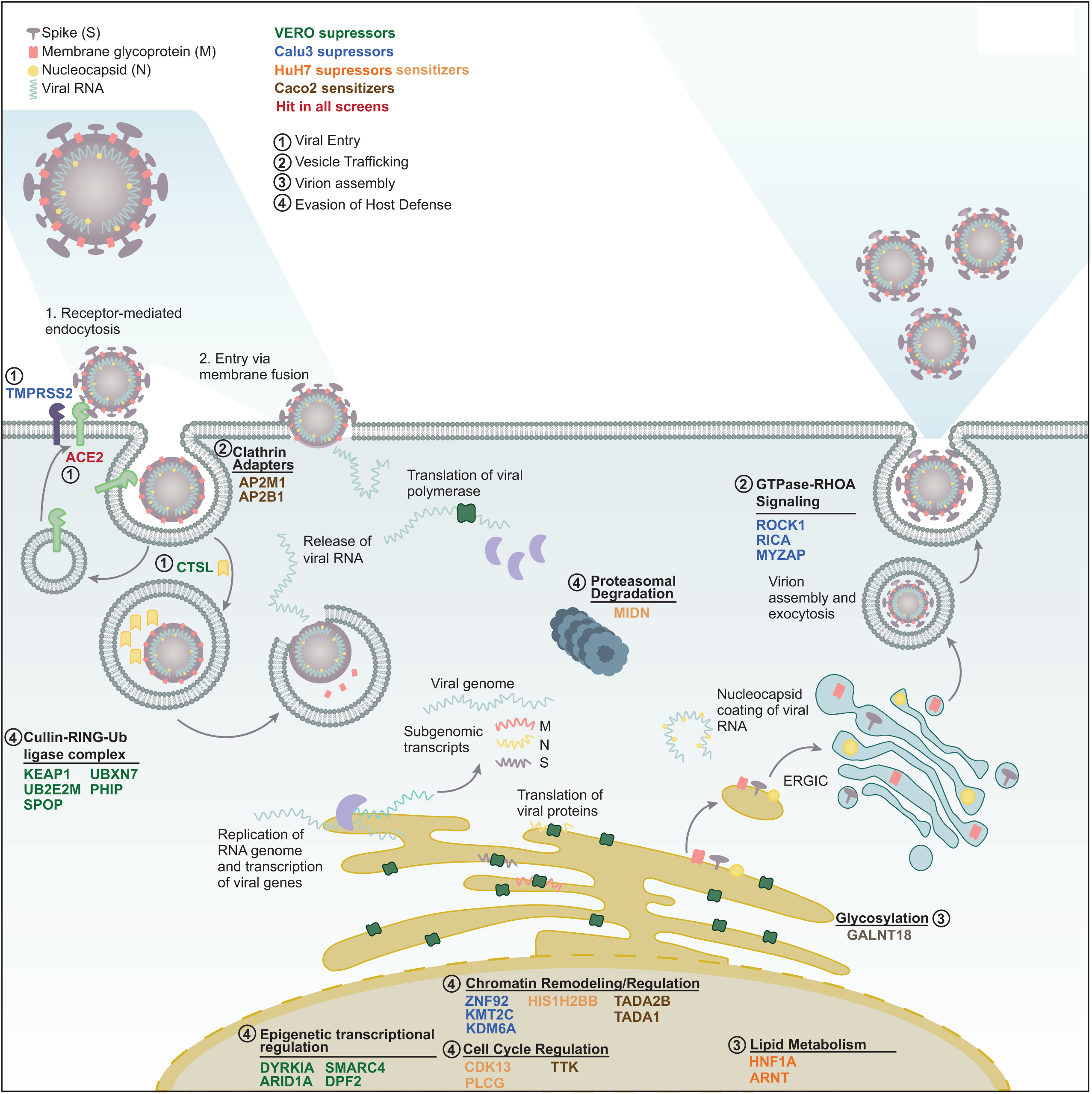
Enriched pro-viral gene hits identify network of host cellular processes that facilitate SARS-CoV-2 infections. Pro-viral (suppressor) genes identified in each cell line are shown in green (vero E6), blue (Calu-3), orange (Huh7) and brown (Caco-2). Four major scenarios were identified for how SARS-CoV-2 potentially utilizes the identified gene hits: 1) viral entry 2) vesicle trafficking 3) virion assembly and 4) evasion of host defences

Huh7 selected for gene hits that could reduce apoptosis including *BMPR1A*, *RGMB*, *SMAD4* are components of TGF-ß/SMAD4 signaling pathway, where RGMB is a GPI anchor member and BMP ligand co-receptor and BMPR1A is a BMP signaling receptor. Modulation of TGF-ß receptors or SMAD factors is one of the main ways virus can interfere with TGF-ß signalling, and SARS-CoV N-proteins have been reported to inhibit formation of SMAD3/4 complexes resulting in reduced apoptosis (Mirzaei and Faghihloo, 2018). Other enriched guides in Huh7 included those targeting *TRIB1*, which encodes an adaptor protein involved in protein degradation and negative regulation of T-cell signalling; *COX2*, a cyclooxygenase proinflammatory gene, that has been shown to be induced by influenza A virus resulting in reduced apoptosis (Dudek et al., 2016); *HNF1A*, which encodes a liver specific transcription factor involved in control of lipid metabolism; and *ARNT*, a gene encoding a transcription factor that dimerizes with AhR (Aryl hydrocarbon receptor) to regulate metabolism by inducing production of cytochrome P4501A1. Both HNF1A and ARNT have been associated with inducing lipid metabolism during viral infections (HBV/HCV respectively), aiding in lipid droplet formation that is required for forming viral particles (Josset et al., 2012; Ohashi et al., 2018). In addition, both co-factors have also been implicated in modulating the immune response by activating NF-kß pathway in response to infections (Josset *et al*., 2012; Lawrence and Vorderstrasse, 2013).

Calu-3 cells, like Huh7 and Vero E6, included genes that were involved in transcriptional regulation (*ZNF292*, *KMT2C*, *KDM6A*). Unique to Calu-3 cells were genes encoding components of the GTPse-RhoA signalling pathways including *ROCK1*, *RICA* and *MYZAP*, which are involved in regulation of cell-to-cell adhesion and phagocytosis, that can aid in spread and entry of SARS-CoV-2. *PRSS8* encodes a serine protease and the interferon-inducible gene *IRF9* were also uniquely enriched in Calu3 cells. Together, these hits in Calu-3 may be involved in viral entry to support virus pathogenesis in Calu3 cells.

Since *ACE2* was not identified in Caco-2 cells and the infection was significantly lower in these cells, there was limited enrichment of pro-viral factors. However, the top genes that were significant included genes that have been implicated as host factors for viral infections; for example *AMBRA1* encodes a regulator of autophagy and has been shown to be degraded by viral proteins HIV and HPV (Antonioli et al., 2020; Nardacci et al., 2014); *PVRL1* encodes a type1 membrane glycoprotein and known entry receptor for HSV, also involved in cell to cell spread (Martinez and Spear, 2001), and *EIF4G1* encodes a component of translation initiation complex which can be recruited by picornavirus IRESs to hijack the host translation machinery (Avanzino et al., 2017).

The top genes (normZ <-3, FDR<0.2) whose perturbation sensitized cells to SARS-CoV-2 cytotoxicity were further evaluated in Huh7 and Caco-2 cells. Since Huh7 or Caco-2 cells did not undergo CPE, negative selections could be performed to capture SARS-CoV-2 evasion genes, or antiviral host factors. Enrichment analysis of the top evasion genes identified processes involved in regulation of cell cycle (*TTK*, *CDK13*, *PLCG*), chromatin regulation (*HIST1H2BB*, *TADA2B*, *TADA1*), glycosylation (*GALNT18*), proteasome degradation (*MIDN*) and motility (*TMOD3*). In Caco-2 cells specifically, genes encoding the AP2-clathrin adaptor complex proteins (*AP2M1*, *AP2B1*) were top evasion genes. AP2 adaptor complexes associate with the plasma membrane and are responsible for trafficking proteins into endosomes for recycling or degradation via lysosomes (Conner and Schmid, 2003). AP2M1 specifically is an intracellular cargo molecule that recognizes the Yxxø sorting motif present in the cytosolic tail of different cargo proteins. This motif is also common in multiple viruses, including SARS-CoV-2, allowing viruses to exploit AP2M1 for internalization from the plasma membrane to intracellular sites (Minakshi and Padhan, 2014; Yuan et al., 2020). Calu-3 knockout screens reported by others (Biering et al., 2021; Rebendenne et al., 2021), also identified members of the clathrin adaptor complex; however, the genes were pro-viral and associated with the AP1 complex, which is found on the trans-Golgi-network and endosomes and is responsible for trafficking proteins between the Golgi and/or endosomes to lysosome (Biering *et al*., 2021; Touz et al., 2004). Many viruses are known to hijack endocytic mechanisms for entry to the cell (Robinson et al., 2018). However, an antiviral role could be the result of two scenarios: trafficking viral proteins to the lysosome for degradation (Park and Guo, 2014), or regulation of signalling pathways following activation by endocytosis of cytokine receptors (Moore et al., 2020; Zanin et al., 2020).

### Comparative analyses of genetic screens confirm little overlap across models

We compared the top 25 resistance gene hits from each screen, including those in previously reported screens in Vero E6(Wei *et al*., 2021), A549^+ACE2^ (Daniloski *et al*., 2021), Huh7.5.1 harbouring ACE2-TMPRSS2 (Wang *et al*., 2021); native Huh7.5 (Schneider *et al*., 2021) and Calu-3 cells (Biering, 2021 #1513). As shown by the Upset plot in Figure 5a, *ACE2* was the only gene that overlapped across all cell types. Not surprisingly, it is more likely for screens performed in the same cell line to have overlapping hits, as we observed when comparing our findings to previously reported Vero E6 and Huh7.5 studies. For example, our Vero E6 screen identified six factors in common with those reported by Wilen et *al.*, including SWI/SNF complex genes *ARID1A*, *SMARCA4* and *SMARCC1*, which were extensively validated (Wei *et al*., 2021). Apart from *ACE2, TMPRSS2* and *CTSL*, 10 genes were individually common between two or three cell lines including *KMT2C, ROCK1, CDH1, EPT1, AMBRA1, HNF1B, EP300,* and *DYRK1A,* and these usually overlapped between Huh7 or Vero E6 (Figure 5a).

**Figure 5.**
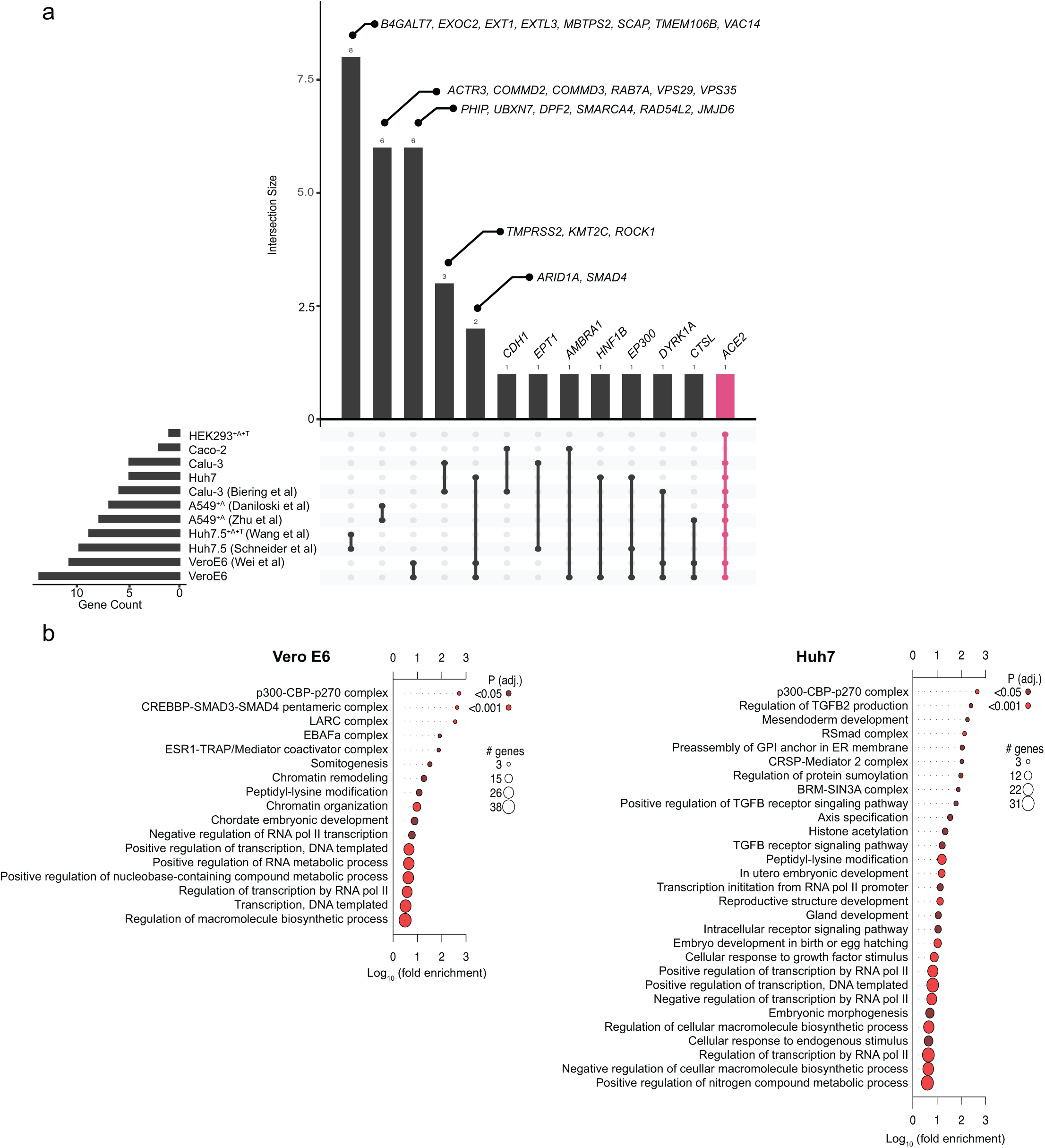
Comparison of overlap between different SARS-CoV-2 genome-wide CRISPR screens A) Upset plot showing overlap of the top 25 hits from this study and previously reported screens B) Enriched GO biological processes for the top 100 resistance hits in Vero E6 and Huh7 cells

Gene Ontology enrichment analysis of the top 100 resistance genes showed that only Vero E6 and Huh7 screens had functionally coherent subsets of genes involved in biological processes involved in SARS-CoV-2 cytotoxicity (Figure 5b). Although the individual gene hits were not the same, similar biological processes were identified, including positive regulation of transcription, RNA metabolism and biosynthesis, and the p300-CBP-p270-SWI/SNF complex. While no functional enrichment groupings were identified for hits selected from Calu-3, Caco-2 or HEK293^+A+T^ screens. Overall, the results show that host genes required for RNA replication or transcription were enriched in two separate cell lines (Vero E6 and Huh7), as well as from the overlaps identified between all screens reported to date; all other enrichments observed for pro-viral factors in previous studies were unique to the screen. The lack of significant overlap between screens in different host cell lines even within the same tissue type, aside from the obligate need for *ACE2*, suggests that cell line is the dominant factor, and that SARS-CoV-2 can utilize an assortment of cellular pathways to create an environment that is favorable for virus replication, which remains to be fully understood.

### Genetic screen validation

Since SARS-CoV-2 was most pathogenic in our Calu-3 screen, we focused on this cell line to validate top gene hits whose knockout elicited resistance to infection. Nine genes out of the top ranked hits were selected for validation including *PRSS8*, *IRF9*, *MYZAP*, *ABCF1*, *PLN*, *HDAC6*, *FAM122B*, *KDM6A*, and *KMT2C*. These genes were selected based on their consistency with known virus-host functions and whether they were novel hits that have not been reported. We also included sgRNAs targeting known factors TMPRSS2 and ACE2 as infection controls. Polyclonal CRISPR knockouts were generated for each gene by transducing gRNAs individually with Cas9 ribonucleoproteins (RNPs), which resulted in high indel frequencies (Supplemental Table 5). Cell viability and viral infection were assessed through detection of N-protein by immunofluorescence to determine whether knockout of the selected genes resulted in resistance to SARS-CoV-2 infection (Figures 6 a,b). As expected, loss of well-established host factors *ACE2* or *TMPRSS2* protected against SARS-CoV-2 infection and virus induced CPE. Loss of chromatin modifying genes *KDM6A* and *KMT2C* significantly reduced SARS-CoV-2 infection and CPE, validating these genes are required for robust SARS-CoV-2 infection. Notably, *KMT2C* is a component of the MLL2/3 complex (also known as the ASCOM complex), which also includes the product of the *KDM6A* gene, and can promote chromatin remodelling by activation via the SWI/SNF complex (Schulz et al., 2019). *ACE2* expression in all nine knockout cell lines was evaluated to determine if the hits selected impacted *ACE2* levels, resulting in their selection as resistance hits (Figures 6 c,d). No changes in the surface expression of ACE2 or total protein was detected, confirming that loss of these genes did not impact ACE2 expression or localization.

**Figure 6.**
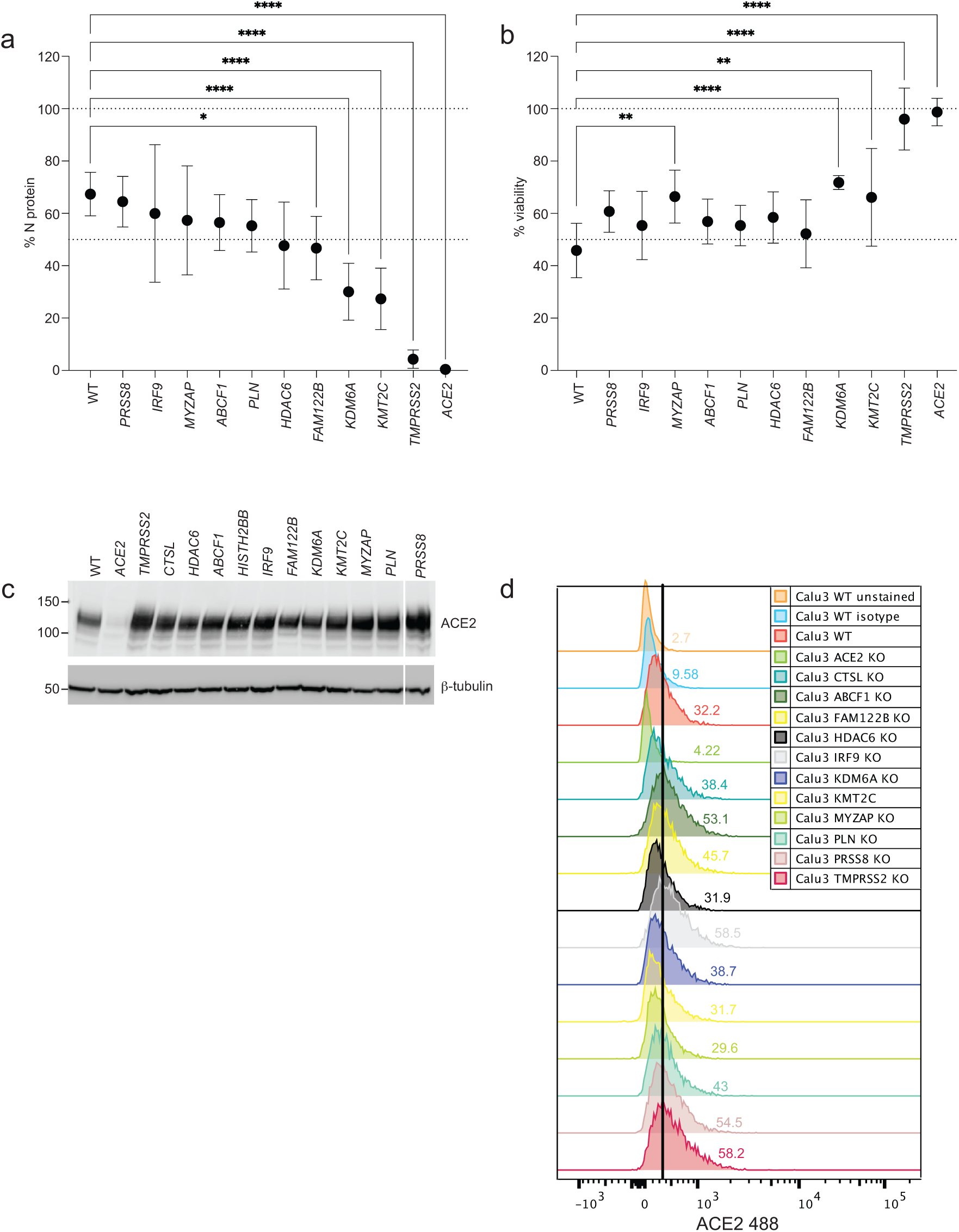
Validation of resistance gene hits sgRNA targeting individual genes were introduced into Calu-3 cells, which were then challenged with SARS-CoV-2 at MOI 1. A) SARS-CoV-2 N-protein and B) cell viability was measured at 48 hpi and compared to WT. ACE2 expression was measured by C) western blot and D) flow cytometry.

### Impact of protease rewiring on virus infectivity

TMPRSS2 and CTSL activity has been reported to be important for viral transmissibility (Hoffmann *et al*., 2020; Zhao *et al*., 2021). After binding to the ACE2 receptor, the SARS-CoV-2 S-protein uses TMPRSS2 on the cell surface or the lysosomal endopeptidase CTSL following clathrin-mediated endocytosis (Hoffmann *et al*., 2020; Zhao *et al*., 2021). Our screens revealed that the protease used for SARS-CoV-2 entry was cell type dependant. In Vero E6 cells, in which TMPRSS2 expression is low, *CTSL* guides were enriched. CTSL was also reported in both Huh7.5 and A549^ACE2+^screens; however, it was not a prominent scoring gene (Daniloski *et al*., 2021; Schneider *et al*., 2021). These data suggest in the absence of TMPRSS2, SARS-CoV-2 can infect cells using an alternative late entry pathway, via endocytosis. Notably, *TMPRSS2* was not a top hit in HEK293^+A+T^, in which infection was thought to be aided by exogenous expression of TMPRSS2 (Figure 3c). However, Calu-3 cells, where CTSL RNA expression was higher than TMPRSS2 (Supplemental Figure 5) were dependent on TMPRSS2 for entry.

We sought to investigate whether proteases utilized in different cell lines are mutually exclusive, leading to a different infection route, or whether it simply depends on the protease expression profile in each cell line. In addition to Vero E6 and Calu-3 cells, we included a bladder cancer cell line UM-UC-4. UM-UC-4 displayed the highest RNA expression of *ACE2, TMPRSS2 and CTSL* in the Cancer Cell Line Encyclopedia (Barretina et al., 2012), and as result was highly susceptible to SARS-CoV-2 infection (Supplemental Figure 6). To determine whether the selectivity of S-protein priming by CTSL or TMPRSS2 can be rewired, TMPRSS2 and CTSL genes were either disrupted using CRISPR or stably overexpressed in Vero, Calu-3, and UM-UC-4 cells. In Calu-3 cells, which were previously TMPRSS2 dependent, became resistant to SARS-CoV-2 infection when TMRPSS2 was knocked out (Figure 7a). Overexpression of CTSL in the presence or absence of TMPRSS2 had no impact on SARS-CoV-2 infection in these cells (Figure 7a). Similar results were observed in Vero E6 cells, in which SARS-CoV-2 entry was CTSL dependent. Overexpression of TMPRSS2 in the presence or absence of CTSL did not result in increased viral transmission (Figure 7b). Interestingly, individual knockout of either TMPRSS2 or CTSL in UM-UC-4 cells did not decrease virus infection (Figure 7c). TMPRSS2-CTSL double knockout in UM-UC-4 cells also did not change the course of SARS-CoV-2 infection compared to WT (Figure 7c), suggesting another cell type that was not dependent on either protease for entry. It is unclear whether another host protease facilitated the cleavage function in the absence of TMPRSS2 and CTSL, or whether cell entry was carried out independently of spike cleavage.

**Figure 7.**
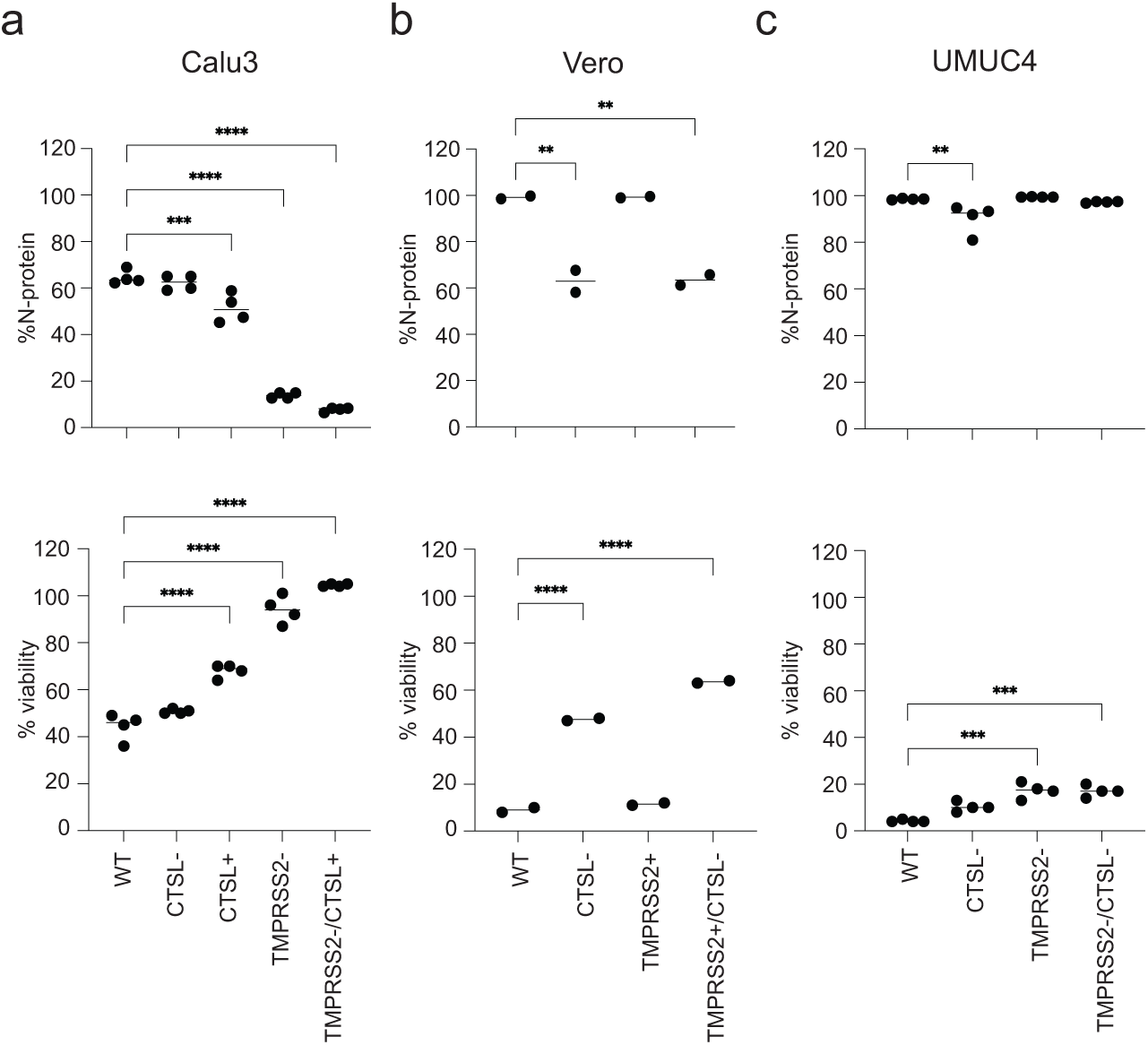
TMPRSS2 and CTSL rewiring for S-protein priming Cell type-specific dependence on TMPRSS2 or CTSL for host cell entry was evaluated by overexpression and knockout of TMPRSS2 in CTSL-dependent cell types and CTSL in TMPRSS2-dependent cell types. Modified cells were challenged with SARS-CoV-2 at MOI 1, and 48 hpi changes to virus transmissibility was measured by immunostaining for SARS-CoV-2 N-protein and DAPI for cell viability in A) Calu-3, B) Vero E6 and C) UM-UC-4 cells.

### Global transcriptional changes to host cells in response to SARS-CoV-2

The global transcriptional changes in host cells in response to SARS-CoV-2 infection was also investigated. Calu-3 and Huh7 cells, cell lines representative of a CPE and non-CPE outcome of SARS-CoV-2 infection respectively, were infected with an MOI∼1. Poly A+ RNA was extracted from infected cells at 0, 12 and 24 hpi, and alterations in mRNA levels, patterns of mRNA splicing and mRNA 3’-end formation (alternative polyadenylation, APA) were assessed (Supplemental Figure 7, Supplemental Table 6). In Huh7 cells, 274 genes were differentially expressed relative to mock samples at 12 hpi. Enriched among genes with reduced expression were transcription factor genes, of which six function in immune response pathways or are known to be differentially expressed upon viral infection (*PPARA*, *FOXO1*, *FOXO3*, *JUN*, *JUNB* and *BLC6)*. An unrelated set of 92 genes was differentially expressed at 12 hpi and 58 genes at 24 hpi. Genes induced at 24 hpi were preferentially involved with type I interferon production and signaling, consistent with a recent report (Banerjee et al., 2021b). Only minor changes to alternative splicing or APA during virus infection was observed (Supplemental Figure 7).

## DISCUSSION

Here we report genome-wide loss of function CRISPR screens in multiple host cell lines including human lung cells (Calu-3), intestinal cells (Caco-2), liver cells (Huh7), embryonic kidney cells (HEK293^+A+T^) and African green monkey kidney cells (Vero E6), to identify host genes required for SARS-CoV-2 infection and cytotoxicity. ACE2 was the only gene that was common across all cell lines in this study and ranked higher as a hit depending on the extent of the cytopathic effect (CPE). For example, *ACE2* was the top hit in Vero E6, Calu-3 and HEK293 cells, while in Huh7 cells that did not exhibit CPE from SARS-CoV-2 infection, *ACE2* was a modest hit, and was not observed in the Caco-2 screen, where infection was low with no CPE. Apart from ACE2, each cell line identified its own unique set of gene hits. The host factors identified from each screen, as summarized in Figure 4, were also involved with other viral pathogens, and speculated to be involved in mechanisms which viruses can evade host cytoprotective processes and provide a permissive environment for virus pathogenesis. The gene hits identified in Figure 4 imply, that SARS-CoV-2 interferes with levels of factors involved in host signalling pathways such as TGF-ß and NF-kB (signalling complexes like Cullin Ring ligases, SMAD4) and shutting down host translation machinery to inhibit gene expression of proinflammatory genes (transcription factors, chromatin remodeling factors). Additionally, genes involved in lipid metabolism, vesicle trafficking and phagocytosis (GTPase RhoA signalling and Clathrin adaptor proteins) were also identified and thought to aid in viral particle formation and viral trafficking. Chromatin modifying genes *KMT2C* and *KDM6A* were the only novel hits that confidently validated to resist SARS-CoV-2 infection. Factors involved in chromatin modification and regulation of transcription, suggest that SARS-CoV-2 may hijack a host of chromatin regulators to reprogram the host transcriptome for successful infection and provide a permissive environment for virus pathogenesis (Wei *et al*., 2021) (Schneider *et al*., 2021). The effects of SARS-CoV-2 infection on the human transcriptome in Calu-3 and Huh-7 infected cells was also analyzed to determine what cellular processes SARS-CoV-2 may co-opt or perturb to satisfy their requirements for growth. Only slight changes in immune pathways were observed at 24 hr. Coronaviruses, such as those that cause SARS and MERS have evolved multiple proteins that can inhibit type I IFN expression (Chen et al., 2014; Lu et al., 2011; Lui et al., 2016; Niemeyer et al., 2020; Schroeder et al., 2021; Siu et al., 2014a; Siu et al., 2014b). Correspondingly, Nabeel-Shah et al, recently showed that SARS-CoV-2 N-protein regulates the host gene expression by attenuating stress granules like G3BPs and binding directly to target mRNAS, countering host responses to viral infection (Nabeel-Shah et al., 2020).

Multiple studies have been reported earlier this year assessing SARS-CoV-2 host factors. Majority of cell lines screened required exogenous expression of ACE2 +/- TMPRSS2 to aid SARS-CoV-2 infection. Similar to the observations in this study, little overlap of gene hits was observed between our screens and reported screens other than ACE2. Additionally, most of the gene functions reported converged back to regulation of *ACE2* host dependency factor, rather than direct viral interactions. As suggested in Figure 5, screens done in identical cell lines overlapped more than between different cell types. A549 screens enriched for processes regulating endosomal processes, such as endosomal acidification, sorting and recycling, linking to the endosomal entry pathway, the primary pathway used by A549 cells for virus transmission. Perturbation of the genes enriched in in these pathways, also perturbed ACE2 cell surface expression (Zhu *et al*., 2021). Huh7 [Hu7.5, 7.5.5] derived liver cells, identified different host factors, but also were linked to receptor usage. These enrichments included GAG biosynthesis, cholesterol homeostasis and RAB GTPases that were shown to influence ACE2 trafficking to the cell surface (Daniloski *et al*., 2021). The hits identified in our Vero E6 screen overlapped with another screen performed by Wei et al(Wei *et al*., 2021) in the same Vero E6 cell background. As other reports have shown, hits have been identified as ACE2 regulators, like RAB7a in A549 cells (Daniloski *et al*., 2021) and HMGB1 in Vero screens (Wei *et al*., 2021). ACE2 levels were not perturbed in our validation studies indicating that the hits selected in our screens, such as chromatin modifiers KMT2C and KDM6A did not converge at regulation of ACE2, Figure 6.

Despite the intense efforts, further characterization work is required to understand the impact of the identified host factors across all screens reported on regulation of SARS-CoV-2 replication. It is clear, however that cellular entry of SARS-CoV-2 depends on the binding of spike (S) protein to ACE2 and subsequent S priming by cellular proteases TMPRSS2 and cathepsin B/L activity as a substitute. The binding affinity of the S-protein and ACE2 was found to be a major determinant of SARS-CoV-2 replication rate and disease severity(Hoffmann *et al*., 2020). Here, ACE2 was the strongest hit and the limiting factor for viral entry at the initial stage of infection, as cell lines that expressed higher levels of ACE2 could be infected with greater efficiency in cell culture. UM-UC-4 cells which expressed the highest levels of ACE2 according to the RNA seq datasets(Barretina *et al*., 2012), were extremely permissive to infection with cells staining 100% positive for SARS-CoV-2 N-protein within in 24h of infection, with high levels of CPE. We identified for the first time alternative human cell lines, such as UM-UC-4 that are susceptible to SARS-CoV-2 infection *in vitro*. In addition, cell types of various tissue origin were shown to be permissive to SARS-CoV-2 infection as long as ACE2 was expressed. The extent of infection correlated to expression of ACE2, as CPE was only observed in Calu-3, UM-UC-4 and Vero E6 that expressed higher levels, and HEK293 cells that were aided by exogenous expression of ACE2. The pathophysiology of SARS-CoV-2 appears to cause multi-organ damage in the clinic (Puelles et al., 2020). ACE2 receptors have been shown to be enriched in alveolar epithelial type II cells of lung tissues, as well as extrapulmonary tissues such as the heart, endothelium, kidneys, and intestines, substantiating the potential for direct viral infection in these tissues contributing to the damage presented clinically (Zaim et al., 2020).

We found TMPRSS2 only in Calu-3 infected cells, while CTSL was only selected in Vero E6 cells (Matsuyama et al., 2020). CTSL was also reported by other groups in Huh7.5 liver and A549^+ACE2^ lung cell screens; however it was not ranked as a top hit. Although, TMPRSS2 activity has been implicated to be important for viral transmission (Sungnak et al., 2020), our results show that it was only essential for infection in Calu-3 lung cells, and loss of this gene inhibited infection while loss of CTSL did not. In contrast, in UM-UC-4 cells, loss of both proteases individually or even in combination failed to block viral infection, indicating TMPRSS2 and CTSL are not mutually exclusive. Hoffman et al (Hoffmann *et al*., 2020) showed that SARS-CoV-2 S-protein can use both CatB/L as well as TMPRSS2 for priming in these cell lines. In contrast, in SARS-CoV-1 infection TMPRSS2 activity is essential for viral spread and pathogenesis, whereas CatB/L are dispensable. It is thought that TMPRSS2 accounts for early entry, where adjacent S-protein are processed at the cell membrane. While late entry occurs when S-proteins traffic to late endosomes/lysosomes for processing by endosomal cathepsins (Park et al., 2016). Knowledge of the proteolytic cleavages and host proteases regulating virus infections can be used to predict viral tropism and pathogenesis and reveal potential antiviral strategies. Ideally targeting the entry steps of SARS-CoV-2 could be an attractive approach to block SARS-CoV-2 invasion of cells. However, the results here suggest the role of TMPRSS2 or CTSL is cell type dependent and correspond to the overall differences in the trafficking routes of the virus. Therefore, from this revelation targeting the protease would not be an ideal target for broad spectrum inhibition of SARS-CoV-2.

Alternatively, ACE2, the most prominent pro-viral host target offers broad spectrum inhibition, against infection to different cell types. In addition, it can address mutational changes of concern that have been centered around the S-protein RBD leading to enhanced binding affinity for ACE2, resulting in increased replication and eventually viral escape, comprising efficacy to treatments and vaccines under development (Linsky *et al*., 2020). The S-protein RBD shows remarkable degeneracy at the ACE2 contact positions with many interface mutations being tolerated or even enhancing (Starr *et al*., 2020). It is not surprising then, that our study revealed, ACE2 as the primary determinant required for SARS-CoV-2 infection. ACE2 was the only gene, whose disruption in all host cell types resulted in complete inhibition of SARS-CoV-2 infection. Antiviral treatments are currently centered on targeting the S-protein RBD or other viral proteins. Recent reports suggest soluble ACE2 can be used as a decoy to neutralize infections and has the potential to have higher affinities for S binding that rival monoclonal antibodies (Chan et al., 2020; Linsky *et al*., 2020).

In conclusion, our study provides a comparative overview of CRISPR screens across multiple cell lines. Our findings reveal a disconnection of hits identified between cell lines, that can also be seen in previously reported screens. It has been suggested that discrepancies between screens could be explained by idiosyncrasies of the particular screen or cell type (Brest et al., 2021). Evaluation of the essential genes in the cell lines revealed context-essentials and cell line specific dependencies, such as DNA damage response in VeroE6. Because most overlaps occurred for screens of the same cell line by different groups using different libraries, provides confidence in the technical quality of CRISPR screens. Across these efforts the consensus of hits and enrichment of ACE2, TMPRSS2 or CTSL, suggests these screens enrich for early-stage host factors, which is typical of pooled cell survival screens using highly cytopathic viruses (McDougall *et al*., 2018). Follow-up screens with more indolent versions of the SARS-CoV-2 may identify dependency factors that act at later stages of replication. Nevertheless, the network of host factors that have been identified will be broadly applicable to understanding the impact of SARS-CoV-2 on human cells and facilitate the development of host-directed therapies. As it stands, the consensus view is that the virus that causes COVID-19 will become endemic, but posing less danger over time(Phillips, 2021). The future will depend on a combination of annual vaccines, acquired immunity and new therapeutics.

## MATERIALS AND METHODS

### Cells

Vero-E6 (ATCC), Caco-2 (ATCC), Huh7 (ATCC), HEK293+A+T (ACE2, TMPRSS2 stable overexpression), UM-UC-4 (ECACC) cells were cultured in DMEM (Wisent) with 10% FBS (Gibco) and 1% penicillin-streptomycin (Gibco). Calu-3, and UM-UC-4 cells were cultured in EMEM (Wisent) with 10% FBS and 1% penicillin-streptomycin. All cell lines were dissociated with trypsin-EDTA (Gibco) and maintained at 37°C and 5% CO_2_. Cells were regularly monitored for mycoplasma contamination.

### Virus stocks

SARS-CoV-2/SB3 isolate (Banerjee *et al*., 2020) used here was propagated in Vero E6 cells at 37°C to generate P4 viral stock. Cells were inoculated for 2-3 days at which cytopathic effects are apparent. Supernatants were harvested, centrifuged, filtered and stored at -80°C. Viral titers of collected supernatants (50% tissue culture infectious dose, TCID50/mL) were determined according to the Spearman and Karber method (Hamilton et al., 1977) as outlined previously(Banerjee *et al*., 2020). All experiments with SARS-CoV-2/SB3 were performed in containment level 3 (CL3) facility.

### SARS-CoV-2 virus infections

Host target cells were seeded overnight to achieve 90% confluence at the time of infection. Monolayers were infected with SARS-CoV-2/SB3 at indicated MOIs for 1 hour at 37°C in serum-free DMEM or EMEM. After 1 hour, the infection inoculum was removed, and cells were washed and replenished with regular cell culture media depending on the cell type. 24-120 hours post infection, supernatants were harvested and stored at -80 and cells fixed with 4% neutral buffered formalin (NBF) for 1 hour at room temperature for further analysis.

### Immunofluorescence and confocal microscopy

To detect SARS-CoV-2 N-protein expression, cell cultures were seeded and infected at the desired MOI in black 96-well PerkinElmer CellCarrier ultra imaging plates (Perkin Elmer). Infected cells were fixed with 10% neutral buffered formalin (NBF) . After fixation, cells were permeabilized with 0.1% Triton X-100 for 10 minutes at room temperature, washed with PBS and blocked for 1 hour in solution containing PBS, 2% BSA and 2% FBS. After blocking, cell were stained overnight at 4°C with SARS-CoV-2 N-antibody (Genscript) at a dilution of 1:500 in PBS + 1% BSA+ 1%FBS. The next day, cells were washed with PBS and incubated with secondary antibody, anti-human FITC-IgG (Fc specific) (ThermoFisher Scientific) at a dilution of 1:400 and for 45 minutes at room temperature. F-actin was visualized in the same samples by incubation with phalloidin-568 (ThermoFisher Scientific). Finally, cells were washed with PBS, and counterstained with Hoechst 33258. Cells were imaged on the Opera Phenix QHES (PerkinElmer) automated high-content screening system. Percentage of cells infected were determined using Harmony^TM^ high content imaging and analysis software by comparing cells stained positive with anti-N antibody versus Hoechst actin-stained cells.

### Western blot analysis

Cells were lysed in RIPA buffer (50mM Tris pH7.5, 150mM NaCl, 1mM EDTA, 1mM EGTA, 0.1% SDS, 0.5% sodium deoxycholate, 1% Triton X-100, 1mM PMSF supplemented with a protease inhibitor cocktail) and centrifuged 14,000 rpm for 15 minutes at 4℃. Cell lysates were mixed with BOLT LDS sample buffer supplemented with a reducing agent (Life Technologies) and boiled for 5 minutes. Next, 10-30 ug of protein was resolved on 4-12% Bis-Tris gels (Life Technologies) and transferred to a nitrocellulose membrane using iBLOT2 with the compatible NC midi stacks (Life Technologies) at 20V for 1 minute, 23V for 4 minutes and 25V for 2 minutes. Membranes were blocked with 3% BSA (BioShop) in 1X PBS prior to antibody decoration. Subsequently proteins were detected using anti-ACE2 (R&D AF933, RRID:AB_439702), anti-TMPRSS2 (Millipore, MABF2158), anti-CTSL (SantaCruz SC-32320 RRID:AB_626811), ß-Tublin (DSHB E7, RRID:AB_528499), anti-HSP90 (SantaCruz, SC-13119, RRID:AB_675659) or anti-FLAG (Sigma, A8592, RRID:AB_439702) antibodies with an appropriate HRP conjugated secondary antibody, anti-mouse-IgG (Cell Signaling Technologies 7076, RRID:AB_330924), anti-rabbit-IgG (Cell Signaling Technologies 7074, RRID:AB_2099233) or anti-goat IgG (Jackson ImmunoResearch 805035180, RRID: AB_2340874). Proteins were visualized on Microchemi 4.2 (DNR Bio Imaging Systems) using Immobilion Western ECL reagents (Millipore).

### ACE2 flow cytometry analysis

For ACE2 labeling, cells were harvested with 5mM EDTA, washed once with FACS buffer (1%BSA in PBS and 5mM EDTA) and incubated for 40 minutes in FACS buffer containing 0.75ug of human ACE2-488 conjugated antibody (R&D Systems FAB9332) and 1:1000 7-AAD (ThermoFisher Scientific). Mouse IgG2A-488 (R&D systems, IC003G) was used an isotype control. Flow cytometry was performed using the LSR Fortessa X20 (BD Biosciences) and analyzed with FlowJo v10 software.

### Genome-wide CRISPR screens

The human Toronto knockout v3 (TKOv3) genome-scale CRISPR library (Addgene #90294) was used to perform pooled CRISPR knockout screens in Vero E6, Caco-2, Huh-7, Calu-3 and HEK293^+A+T^ cells(Hart *et al*., 2017). Host cells were transduced with TKOv3 lentivirus at an MOI of 0.3, such that each sgRNA was represented in about 200 cells. 24 hours after, TKOv3 transduced cells were selected with puromycin (3-12ug/ml) for 48 hours. After 48 hours, cells were harvested, pooled and split into three replicates of at least 1.5×10^7 cells (minimum 200-fold coverage TKOv3 library) and passaged every 3-6 days depending on the doubling time of the host cell line, maintaining coverage at 200-fold. 30×10^6 cells were collected for genomic DNA extraction at T0 and at every passage. After two cell passages transduced cells were infected with SARS-CoV-2 at the determined MOI for each host cell. For cytopathic screens (Vero-E6, Calu-3, HEK293^+A+T^), 2.0×10^7 cells per replicate was infected with SARS-CoV-2 at the indicated MOI. Virus induced CPE was apparent 2-3 days after infection, after which surviving cells were maintained in standard growth media until confluence. SARS-CoV-2 treated cells were collected for genomic DNA extraction and passaged for repeat infection with SARS-CoV-2 until no CPE was observed. For non-cytopathic drop out screens (Huh-7, Caco-2), SARS-CoV-2 infected cells were passaged every 3-4 days, at which cell were re-infected with SARS-CoV-2 for 2-3 rounds. For all screens mock infected cells were passaged and cell pellets collected for genomic DNA every 3-6 days until completion to serve as a reference for sgRNA analysis.

Genomic DNA from all screens was extracted from cell pellets using the Wizard Genomic DNA Purification kit (Promega). Sequencing libraries were prepared from 100 μg of gDNA for control sample and non-cytopathic screen samples or 10 μg of gDNA for cytopathic screen samples via a 2-step nested PCR using primers that include Illumina TruSeq adapters with i5 and i7 indices. Barcoded libraries were gel purified using PureLink Quick Gel Extraction kit (ThermoFisher) and sequenced on an Illumina HiSeq2500 using single-read sequencing and were completed with standard primers for dual indexing with HiSeq SBS Kit v4 reagents as described in (Aregger et al., 2020).

### Screen analysis

Demultiplexed FASTQ files were first trimmed by locating constant sequence anchors and extracting 20-bp guide sequences preceding the anchor sequence. Pre-processed paired-end reads were aligned to TKOv3 reference library sequences using Bowtie (v0.12.8) allowing up to two mismatches and one exact alignment (specific parameters: -v2 -m1 -p4 --sam-nohead). Successfully aligned reads were counted and combined into a matrix with guide annotations. Read counts for all samples were normalized by dividing each read count by the sum of all read counts in the sample then multiplying by the expected read number (10 million). The abundance (fold-change) of each guide was determined for each starting and selected population. The calculated fold-changes were then used to generate normZ scores using drugZ (version1.1.0.2-) to determine enrichment and depletion scores for each gene(Colic *et al*., 2019).

### RNA sequencing

Total cellular RNA was extracted using the RNeasy Kit (Qiagen) according to the manufacturer’s instructions. Samples were submitted for mRNA-Seq at the Donnelly Sequencing Centre at the University of Toronto (http://ccbr.utoronto.ca/donnelly-sequencing-centre). RNA-seq libraries were sequenced on an Illumina NovaSeq6000 platform using an S2 flowcell at 2×151-bp read lengths. Gene expression changes were analyzed by pseudo-aligning pre-trimmed reads to GENCODE v29 transcripts using Salmon v0.14.1 #(Patro R et al., Nat Methods 2017)# and aggregated per gene using the R package tximport #(Soneson C et al., F1000Research 2015)#. Differential expression was then assessed using the classic mode (exactTest) edgeR #((McCarthy et al., 2012))#, and genes with an FDR < 0.05 and a greater than 2-fold change were considered differential. For each comparison, only genes expressed at a minimum of 5 RPKM in one or both conditions were considered. Alternative splicing was analyzed using Vast-tools v2.2.2 (Tapial et al., 2017) in combination with the VastDB Hs2 library released on Dec. 20, 2019 (https://github.com/vastgroup/vast-tools). Changes were considered significant if they were greater than 10 dPSI/dPIR and the expected minimum change was different from zero at p > 0.95 according to vast-tools’ *diff* module. Events were filtered requiring a minimum of 10 reads per event and a balance score (quality score 4) of ‘OK’,‘B1’,‘B2’, ‘Bl’ or ‘Bn’ for alternative exons or > 0.05 for intron retention events in at least 2 of 3 replicates. Alternative cleavage and polyadenylation analysis was performed using QAPA with default settings, essentially as previously described (Ha et al., 2018). QAPA builds the reference library of 3′ UTRs from all annotated protein-coding genes using GENCODE basic gene annotation (v19) for humans (hg19), supplemented by experimentally defined polyA sites archived in the PolyAsite database (Gruber et al., 2016). To quantify the poly(A) site usage (PAU), the percentage of expression of a single 3’ UTR over the total expression level of all 3’ UTRs for a given gene was calculated. Lengthening events were defined as those with proximal PAU group difference between two conditions (PAU_Group_diff) < −20% and shortening events with proximal PAU_Group_diff > 20%.

### Gene set enrichment and network analysis

Analysis of enriched gene functions was performed as ordered queries with g:Profiler ###Ref: https://doi.org/10.1093/nar/gkz369 ### using the TKOv3 library as a custom background. The ‘biological process’ domain of Gene Ontology as well as CORUM were selected as annotation sources, and only terms with at least five and at most 1000 members as well as an intersection size with the query of at least 3 genes were considered. For plotting purposes, only terms with at least 3-fold enrichment with respect to the background were used, and when terms mutually overlapped by at least 70% genes present in the input, only the most enriched term was retained.

GO enrichment analysis for differentially expressed genes and genes containing alternative splicing changes was performed with FuncAssociate 3.0 #(Berritz et al., Bioinformatics 2009)#, requiring at minimum a 5-fold enrichment and excluding categories of more than 1000 genes. Terms with a mutual gene overlap of greater then 50% were merged. Gene set analysis for APA changes were performed using the gProfiler R package (v0.7.0) using the GO-Bioprocess, GO-Molecular Function and Reactome pathway standards (Raudvere et al., 2019). For screens, enrichment analysis was performed for significant hits (|normZ| >3, FDR <0.2).

### Mass spectrometry

#### Protein isolation

Proteins were isolated from Trizol following RNA isolation. Briefly, DNA was precipitated by the addition of 0.3 ml of 100% EtOH and discarded. 1.5 ml of isopropanol was added to the remaining mixture to precipitate the proteins. After 30 min incubation at RT (room temperature), the samples were centrifuged for 10 min at 14000 rpm at 4C. The pellets were washed twice with 95% EtOH. The pellets were dried briefly to evaporate ethanol and resuspended in 8M Urea, 50mM Tris, ph=7.9. The pellets were left overnight at 4C to ensure efficient solubilisation. The next day, 8M urea was diluted in half with 50mM ammonium bicarbonate. The proteins were then precipitated using Proteoextract protein precipitation kit (Calbiochem; cat # 539180) according to manufacturer’s instructions. The precipitated proteins were resuspended in 50 mM ammonium bicarbonate and subjected to trypsin digestion.

#### Trypsin digestion

Each sample was resuspended in 88 uL of 50mM NH_4_HCO_3_, reduced with 8mM DTT for one hour at room temperature, alkylated with 500mM iodoacetamide for 45 min in the dark room, and digested with 1μg of trypsin overnight at 37°C. Samples were desalted using ZipTip Pipette tips (EMD Millipore) using manufacturer protocol and dried.

#### Data Acquisition

Peptides were reconstituted in 20μl of 1% formic acid and 5μl was loaded onto the column. Peptides were separated on a reverse phase Acclaim PepMap trap column and EASY-Spray PepMap analytical column using the EASY-nLC 1200 system (Proxeon). The organic gradient was driven by the EASY-nLC 1200 system using buffers A and B. Buffer A contained 0.1% formic acid (in water), and buffer B contained 80% acetonitrile with 0.1% formic acid. 90 min gradient was utilized at a flow rate of 250 nL/min, with a gradient of 0% to 6% buffer B in 1 min, followed by 6% to 30% buffer B in 75 min, 30% - 100% in 4 min, and 100% buffer B for 10 min. Eluted peptides were directly sprayed into a Q Exactive HF mass spectrometer (ThermoFisher Scientific) with collision induced dissociation (CID) using a nanospray ion source (Proxeon). The full MS scan ranged from 300 – 1650 m/z and was followed by data-dependent MS/MS scan of the 20 most intense ions. The resolutions of the full MS and MS/MS spectra were 60,000 and 15,000, respectively. Data-dependent mode was used for MS data acquisition with target values of 3E+06 and 1E+05 for MS and MS/MS scans, respectively. All data were recorded with Xcalibur software (ThermoFisher Scientific).

#### Data Analysis

AP-MS datasets were searched with Maxquant (v.1.6.6.0)(Tyanova et al., 2016). Human protein reference sequences from the UniProt Swiss-Prot database were downloaded on 18-06-2020. SARS-CoV-2 protein reference sequences (GenBank accession NC_045512.2, isolate=Wuhan-Hu-1) were downloaded from https://www.ncbi.nlm.nih.gov/datasets/coronavirus/proteins on 14-09-2020, supplemented by isolate 2019-nCoV/USA-WA1/2020 (GenBank Accession MN985325) for ORF3b, ORF9b and ORF9c. Spectral counts as well as MS intensities for each identified protein were extracted from Maxquant protein Groups file. MS intensity in each sample was normalized to allow across cell line comparisons. Each intensity value in a sample is multiplied by 1e+12, then divided by the total intensity in the entire sample to obtain the normalized intensity.

### Generation of cell lines

Gene-KO cell lines were generated by electroporation (Neon® Transfection system) of Cas9-RNPs. CRISPR guide RNA sequences (gRNA) from TKOv3 library hits were synthesized by IDT. gRNAs were complexed at a 1:1 molar ratio with ATTO550 labelled tracrRNA (IDT) in TE buffer by heating at 95°C for 5 minutes followed by cooling to room temperature to form crRNA:tracrRNA duplexes. 180 pmol of Alt-R Cas9 enzyme (IDT) was combined with 220 pmol of crRNA:tracrRNA duplex (IDT) at room temperature for 20 minutes to form ribonucleoprotein (RNP) complexes in 10ul total volume with Resuspension buffer R (Neon® Transfection system). Target cells were resuspended in Resuspension Buffer R to a dilution of 5×10^3 cells/ul. 90ul (4.5×10^5) of the cell solution was gently mixed with CRISPR RNP complexes and immediately electroporated according to the manufacturer’s protocols (Neon® Transfection system, Thermo Fisher Scientific) and transferred to a pre-warmed 6-well plates for incubation under standard conditions. To confirm gene knockout, genomic DNA from surviving cells was extracted using Extracta DNA Prep (Quanta Bio), Sanger sequencing was performed across the gRNA target sites following PCR amplification and gene knockouts were confirmed using TIDE (https://tide.nki.nl/) to identify out-of-frame insertion-and-deletion mutations.

To generate protease overexpression lines, Tmprss2-FLAG, FLAG-Tmprss2 or Cathepsin L-FLAG constructs were cloned into pLenti6.2 plasmid with blasticidin resistance marker using gateway cloning system (Invitrogen). Lentiviruses were produced by transfecting HEK-293T cells with pPAX2, pVSVG and pLenti6.2 generated above using Lipofectamine 2000 (Invitrogen). The conditioned media was collected from transfected cultures, filtered through 0.22um filters and applied to Vero E6, Calu-3 and Huh7 cells at 1:10 ratio of conditioned media : fresh media, in the presence of 8ug/ml polybrene (Sigma). Transduced cells were washed 24 hours post-transduction, and selected in blasticidin (Gibco) containing media (5ug/ml for Vero and Calu-3, 2ug/ml for Huh7) for 2 weeks. The expression of the FLAG tagged constructs were verified by Western blotting.

## ACKNOWLEDGEMENTS

We would like to acknowledge Anne-Claude Gingras and Payman Samavarchi-Tehrani for sharing HEK293^+A+T^ cell line and Betty Poon for assistance in the CL3 facility. This work was supported by the University of Toronto COVID-19 Action Initiative Fund to J.M., B.J.B., S.G.O., J.G., K.M., and S.M.. Indirect support was also received from the University of Toronto and the Temerty Foundation to support enhanced capacity and operations of the Toronto Combined Containment Level 3 Facility during the COVID-19 pandemic. This work was also partially supported from a Canadian Institutes for Health Research Project Grant to J.M. (MOP142375). J.M. is a Tier 2 Canada Research Chair in Function Genomics.

## AUTHOR CONTRIBUTIONS

Conceptualization and design of study: K.C., A.G.F., H.L., F.G., P.M., U.B., K.R.B., E.M., J.G., B.J.B., M.T., J.M.; Experimental Investigation: K.C., A.G.F, H.L., F.G., P.M., K.A., S.H., E.M., A.H.Y.T., A.H., P.B., A.A., A.B., N.C.H.; Data Analysis: K.C., A.G.F., H.L., F.G., P.M., E.M., U.B., K.R.B., S.P., A.A., M.T., J.M.; Writing and editing: K.C., P.M., A.G.F., H.L., U.B., K.R.B., M.T., J.M. with input from all other authors; Supervision: K.C., N.C.H., P.M., S.M., K.M., S.G.O., J.G., B.R., B.J.B., M.T., J.M.; Funding acquisition: S.M., K.M., S.G.O., J.G., B.J.B., M.T., J.M.

